# How the Zebra got its Rump Stripes: Salience at Distance and in Motion

**DOI:** 10.1101/2021.04.16.440148

**Authors:** Alex Muhl-Richardson, Maximilian G. Parker, Greg Davis

## Abstract

Zebras’ stripes cannot protect them from predators, Darwin concluded, and current consensus tends to support his view^1,2^. In principle, stripes could support crypsis or aposematism, could dazzle, confuse or disrupt predators’ perception^3–8^, yet no such effects are manifest in predator-prey interactions^9–11^. Instead, narrow stripes covering zebras’ head, neck, limbs and flanks are an effective deterrent to tabanids^12^, vectors for equine disease^13,14^. Accordingly, while other potential benefits, e.g., thermoregulation^15,16^ and intraspecific communication^17^, cannot be excluded, zebra stripes likely evolved primarily to deter parasites^18–20^. Rump stripes, however, do not fit this, or any extant view. Typically horizontal and broader in sub-species with width variation, they are ill-suited to crypsis or parasite-deterrence^12^ and vary with hyaena threat^18^, perhaps shaped by an additional selective pressure. We observed that rump (and rear-flank) stripes remain highly conspicuous when viewed in motion or at distance, while other stripes do not. To study this striking effect, we filtered images of zebra to simulate acuity limitations in lion and hyaena photopic and mesopic vision. For mountain zebra and plains zebra without shadow striping, rump stripes were the most conspicuous image regions according to computational salience models, corroborated by human observers’ judgements of maximally attention-capturing image locations, which were strongly biased toward the rear. By hijacking exogenous attention mechanisms to force predator attention to the rear, salient rump stripes confer benefits to zebra, estimated here in pursuit simulations. Benefits of rump stripe salience may counteract anti-parasite benefits and costs of conspicuity to shape rump and shadow stripe variation.

**Highlights:**

- Zebra stripes likely evolved to deter biting flies, but rump stripes are ill-suited to this.
- Rump-stripes remain highly conspicuous when viewed at distance or in motion.
- Computational models and human observers’ judge rump stripes are most salient stripes.
- Salient rump stripes drive predator attention to rear, hindering capture by predators.
- Observe this striking effect in moving zebra at: viscog.psychol.cam.ac.uk/resources-and-downloads

## 2 Results and Discussion

Figure 1 shows saliency heatmaps for example images from four different striping-patterns in Experiment 1: first column - mountain zebra (Equus zebra zebra, E. z. hartmannae), second column - plains zebra without shadow stripes, ‘plains-’ (Equus quagga borensis, E. q. boehmi, E. q. chapmani), third column - plains zebra with salient shadow stripes, ‘plains+’ (E. q. burchellii), and fourth column - Grevy’s zebra (Equus grevyi). Each heatmap indicates maximally salient regions of each image calculated from observers’ judgments of maximally attention-capturing locations (first and third rows) or an example computational model of saliency (second and fourth rows; Learning Discriminative Subspace [LDS] model). Blurring in images from the static condition illustrates basic filtering to simulate leonine photopic acuity and dichromacy^21^; simulated motion images were additionally manipulated to simulate the effects of horizontal retinal-image blurring when a predator fails perfectly to track lateral image motion at close range (see Methods).

**Figure 1:**
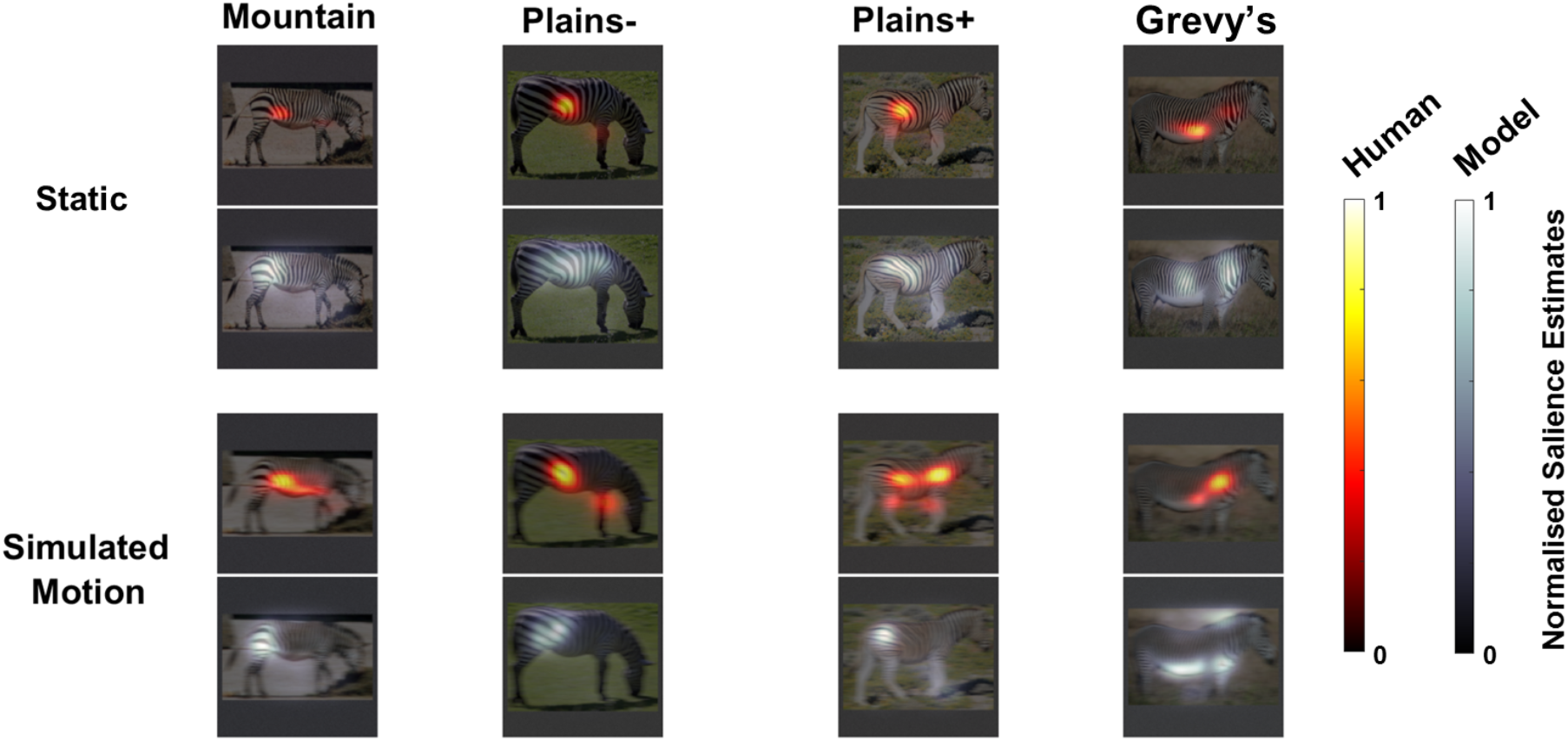
Example salience heatmaps for human observers and Learning Discriminative Subspace model estimates for striping patterns in Experiment 1, heatmap scale shows a normalised salience estimate.

In agreement with our pre-registered predictions for Experiment 1 (AsPredicted.org #52944), the maximally attention-capturing locations selected by observers were subject to an effect of simulated motion, with strong biases to the rear of mountain zebra, *F*(1,23) = 24.51, *p* < .001, *η*^2^_G_ = 0.17, and plains zebra without shadow stripes, *F*(1,23) = 8.23, *p* = .009, *η*^2^_G_ = 0.07 (see Figure 2, first column). Grevy’s zebra, *F*(1,23) = 24.84, *p* < .001, *η*^2^_G_ = 0.20, and plains zebra with shadow stripes, *F*(1,23) = 8.23, *p* = .009, *η*^2^_G_ = 0.07, were also subject to an effect of simulated motion which pushed observers’ judgements rearward relative to static conditions.

**Figure 2:**
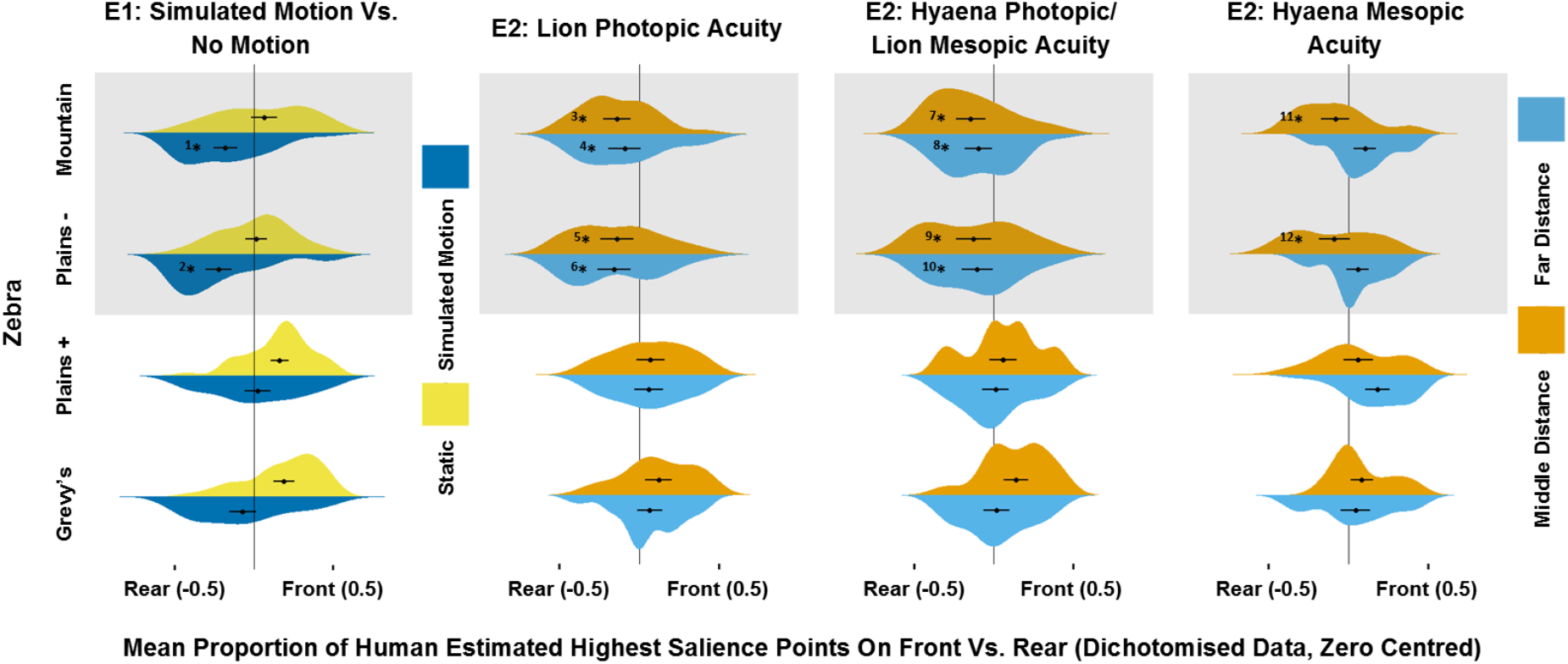
Mean dichotomised proportions of human salience estimates to front and rear of different striping patterns. The first column shows the static and simulated motion conditions in Experiment 1. Under simulated motion, both mountain zebra, 1 - *t*(23) = 3.62, *p* < .001, *d* = 0.74, and plains zebra without shadow stripes (plains -), 2 - *t*(23) = 4.03, *p* < .001, *d* = 0.82, showed salience estimates with a rearward bias. The second, third and fourth columns show the far/middle distances and three acuity levels (lion photopic, hyaena photopic/lion mesopic, hyaena mesopic) in Experiment 2. In line with predictions, mountain zebra and plains zebra without shadow stripes showed salience estimates with a rearward bias in all conditions except when the hyaena mesopic acuity was coupled with far distance, 3 - *t*(27) = 3.29, *p* = .001, *d* = 0.62, 4 - *t*(27) = 1.84, *p* = .038, *d* = 0.34, 5 - *t*(27) = 2.73, *p* = .005, *d* = 0.52, 6 - *t*(27) = 3.07, *p* = .002, *d* = 0.58, 7 - *t*(27) = 3.14, *p* = .002, *d* = 0.59, 8 - *t*(27) = 2.28, *p* = .015, *d* = 0.43, 9 - *t*(27) = 2.31, *p* = .014, *d* = 0.44, 10 - *t*(27) = 2.20, *p* = .018, *d* = 0.42, 11 - *t*(27) = 1.90, *p* = .034, *d* = 0.36, 12 - *t*(27) = 1.86, *p* = .037, *d* = 0.35. Error bars show 95 percent confidence intervals. Note - these data do not reflect how far salience estimates were towards the front or rear, only the proportions that were on the front vs. the rear.

In planned analyses including both simulated motion and static images, an overall rearward bias was detected for plains zebra without shadow stripes, *F*(1,23) = 5.85, *p* = .024, *η*^2^_G_ = 0.14, but not for mountain zebra, *F*(1,23) = 1.64, *p* = .212, reflecting an absence of any bias for static images. Exploratory analyses confirmed effects only in the simulated motion condition (see Figure 2 caption). As expected, no overall bias was evident for Grevy’s zebra, *F*(1,23) = 2.14, *p* = .157, whereas an opposite bias was observed plains zebra with shadow stripes, *F*(1,23) = 6.17, *p* = .021, *η*^2^_G_ = 0.14.

This transformative effect of motion - amplifying the relative salience of broad, horizontal rump-stripes, versus of narrow, often vertical stripes - can readily be observed in footage of moving zebra (viscog.psychol.cam.ac.uk/resources-and-downloads). This effect was weaker in plains zebra with shadow stripes and while simulated motion eliminated the frontward bias evident in the static condition, it did not reverse it. This is consistent with our view that thinner shadow stripes enhance tabanid deterrence^12^ at the cost of reduced rump salience. One unexpected finding was that simulated motion did induce a rearward salience bias in Grevy’s zebra images. Further planned comparisons revealed that both mountain zebra, *F*(1,23) = 15.05, *p* < .001, *η*^2^_G_ = 0.05, and plains zebra without shadow stripes, *F*(1,23) = 40.11, *p* < .001, *η*^2^_G_ = 0.09, yielded greater overall rearward biases than Grevy’s zebra. These analyses did not reveal any significant interactions involving zebra stripe pattern or any other factors, *Fs* <= 1.94.

Points of maximum salience from observers’ judgments and the computational salience model correlated well for simulated motion images (mountain: .419, plains-: .457, plains+: .477, Grevy’s: .378) and static images (mountain: .212, plains-: .363, plains+: .280, Grevy’s: .183). These results suggest that the elements of visual perception captured by the salience model correlated well with the perceptual judgements made by human observers; see Table S2.

Experiment 1 simulated only lions’ perception of moving zebra at close-range. However, to accrue sizeable benefits, rump-stripe saliency must force attention to the rear at greater distances for lions and for spotted hyaenas. As the primary effects of rump salience at these distances cannot depend upon substantial retinal image motion (angular velocities decrease with distance) we simulated perception for static viewing conditions under three levels of visual acuity, simulating leonine photopic vision (high), lion mesopic/hyaena photopic vision (medium) and hyaena mesopic vision (low). We scaled images to produce retinal images of approximately the same size as zebras viewed from between 17.28 m and 24.22 m (middle distance) and between 34.56 m and 48.47 m (far distance).

Consistent with our pre-registered predictions (AsPredicted.org #54384), the maximally attention-capturing locations selected by observers settings for mountain, *F*(1,27) = 4.31, *p* = .048, *η*^2^_G_ = 0.10, and plains zebra without shadow stripes, *F*(1,27) = 4.73, *p* = .039, *η*^2^_G_ = 0.12, were biased toward the rear irrespective of acuity and distance. The predicted exception to this pattern, highlighted by the interaction between acuity and distance for mountain, *F*(2,54) = 5.30, pG-G corrected = .011, *η*^2^_G_ = 0.02, and plains zebra without shadow stripes, *F*(2,54) = 13.37, pG-G corrected < .001, *η*^2^_G_ = 0.05, was at the longest distance and lowest acuity level, at which stripes were barely discernible. Figure 3 shows example saliency heatmaps for these striping patterns in Experiment 2, derived from human observers’ responses (left image in each pair) and a computational salience model (right image in each pair), for the middle distance images. In contrast, plains zebra without shadow stripes, *F*(1,27) = 4.63, *p* = .041, *η*^2^_G_ = 0.10, and Grevy’s zebra, *F*(1,27) = 7.91, *p* = .009, *η*^2^_G_ = 0.13, showed overall frontward biases. As in Experiment 1, there was good general agreement between observers’ judgements and the computational model (mean Pearson’s correlation across simulated acuity levels and viewing distances: mountain: .487, plains-: .525, plains+: .427, Grevy’s: .299; see Table S2).

**Figure 3:**
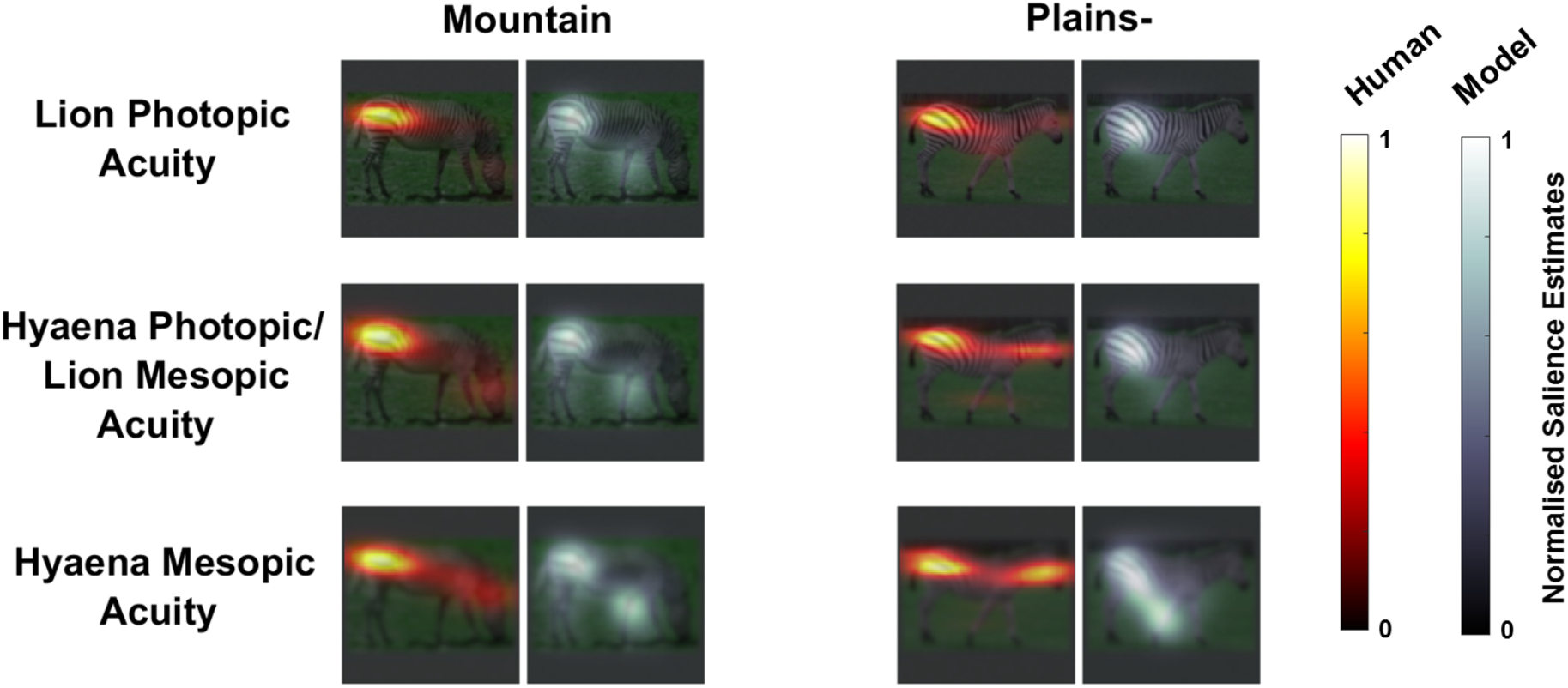
Example salience heatmaps for human observers and Learning Discriminative Subspace model estimates for mountain zebra and plains zebra (without shadow stripes) patterns in Experiment 2, heatmap scale shows a normalised salience estimate.

Human judgments and computational models provide convergent support for our primary claim: that rump stripes’ characteristics make them more salient in most zebra (sub-species of mountain zebra and plains zebra without shadow stripes) than other stripes, when viewed in motion or at distance. We simulated predator acuity for phototopic and mesopic vision to estimate the approximate distances and speeds over which this mechanism may affect predation. However, our claim depends neither on the veracity of estimates, nor on the exact shape of lion and hyaena contrast sensitivity functions. Viewing even standard photographic images and video with (superior) human photopic acuity shows these effects strongly – presumed acuity limits of predators simply alter the ranges at which the effect is strongest. While direct measurement of perceptual salience in lions or spotted hyaenas is not feasible, our results should hold for a broad range of different contrast sensitivity functions.

Using these estimates, we simulated the benefit of forcing predator attention to the rump versus to the head (for a zebra moving predictably and at constant speed orthogonally to a lion’s initial orientation and a lion that accelerates linearly, with a top speed 1.9 times that of the zebra and a maximum pursuit time of 20 seconds^9,22^ see Method for details). This scenario, in which risk to the prey is particularly high, is illustrated in Figure 4 (left panel). In this simulation, if the predator tracks the front of the animal (indicated by red triangles) it catches the zebra, but, if it tracks the zebra’s rear, the predator lags behind. The two middle plots indicate time-to-capture for a broad range of predator velocities (to incorporate diverse estimates of lion and zebra relative velocities). The key panel is on the right – a subtraction of the two middle plots yielding expected increase in time-to-capture as a result of driving attention to the rear. Other than the large black region in the top-left hand corner (slow predators that capture prey in neither case) or starting distances of 10 or less (where the prey is always captured), driving attention to the rear benefits zebra across a broad range of parameter values (by 0-1.5 s, 0-30 m).

**Figure 4:**
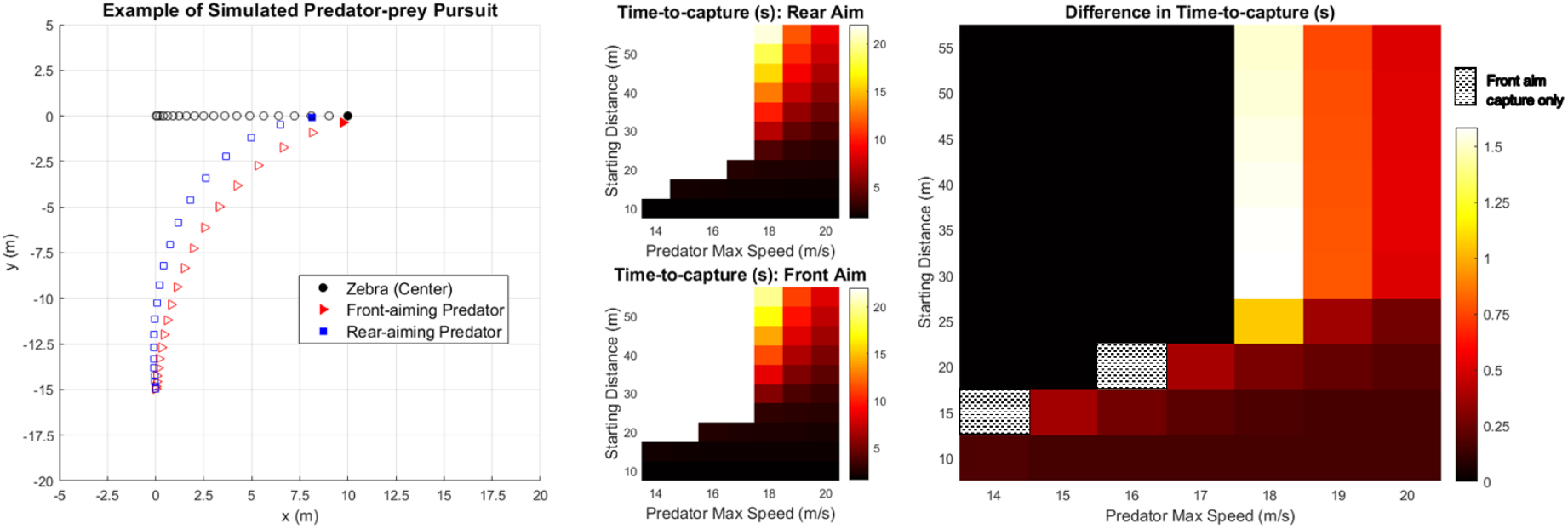
Predator-prey pursuit simulations with time-to-capture estimates for various starting distances and speeds and differences for front-vs rear-aiming predators.

We conclude that, under viewing conditions typical of predator-prey pursuits, rump-stripes of mountain zebra and plains zebra (in the absence of salient shadow stripes), are highly salient, relative to other stripes. We further expect this marked asymmetry to drive predators’ attention to the animal’s rear. Though measuring lion and hyaena visual attention under pursuit conditions is not feasible, mechanisms of visual salience and its exogenous control of visual attention are highly conserved^23^ and would be expected to cue attention in this manner. Further, human observers’ judgements that rump stripes, in images which simulate predators’ limited acuity, captured their attention offers corroborative evidence for this assumption.

In simulation, we established that this mechanism can offer substantial advantages to zebra when pursued by predators. Though the relative importance of speed and acceleration in predator-prey interactions versus rapid turns^22^ remains uncertain, it is clear that speed does play an important role. Ambush predators are thought to hold acceleration rather than speed advantages over their favoured prey species with disadvantages in stamina, while coursing predators rely on stamina and cooperation to bring down larger prey such as zebra^9–11^. For either type of pursuit, a primary requirement for a predator is to make up ground between them and prey before metabolic and thermal-load costs exceed thresholds for termination. This is most easily and safely achieved when the prey’s heading is approximately orthogonal to the predator’s initial heading. Our simulations indicate that, for this illustrative example, driving attention to the rump maximises likelihood of escape, over a range of typical charge distances and relative predator velocities. Additionally, at close quarters, this mechanism might drive attention away from a zebra’s vulnerable head and neck, toward the rump. Patterns of wounding noted by Caro^1^ are potentially consistent with such an effect; careful comparison with patterns from Grevy’s zebra and plains zebra with strong shadow striping, will be required to exclude survivorship effects in those data.

It seems likely, on the basis of these and other findings, that patterns of rump stripes in zebra have been shaped by predation (and flank striping in plains zebra). Though stripes probably evolved primarily to deter biting flies, homogeneous narrow stripes would best serve this purpose. This anti-parasite view, articulated by Harris^19^, Waage^14^, and particularly, Caro^1^ seems the most promising single candidate account, but leaves unexplained a particularly salient feature of zebra stripes – the striking variation in rump stripe width. Our finding here provides an important clue as to the further selective pressures likely responsible for that variation.

Here, to simplify stimulus preparation and observers’ task (indicating the most attention capturing region of an image), we used static stimuli. However, the salient attention cue to the animal’s rear from rump stripes may be enhanced in plains zebra for dynamic stimuli. As is evident in much publicly available footage of zebra, as the hind limbs reach the foremost phase of the stride in locomotion, rump-stripe salience appears maximised as the stripes are horizontal. As rump-stripe orientation changes at other phases of the stride, this modulates the signal in both orientation and salience, exerting a continuous pull on attention.

Our findings also provide a clear mechanism for understanding shadow stripe variation in terms of two selective pressures. In many Burchell’s zebra, salient shadow stripes are often interspersed with rump stripes, or rump stripes can be faint, giving rise to a uniformly lighter pelage. Either of these may deter tabanids but at the cost of reduced antipredator benefits of rump-stripe salience. Accordingly, variation in rump striping pattern may largely reflect competitive interaction of these two selective pressures. Indeed, achieving anti-parasite effects via narrow, salient stripes or more uniform, higher mean reflectance would also explain why leg-stripe salience in Burchell’s zebra is not maximised, though they are likely exposed more than mountain and Grevy’s zebra to horseflies. A two-factor approach to understanding this variation^8^ may help understand while a species level analysis by Caro and colleagues^18^ found that those stripes varied with hyaena prevalence, rather than tabanid exposure, whereas in sub-species analyses trends were clearer for tabanids than hyaenas.

We do not address in detail, here, other previous work on zebra stripes’ potential antipredator functions (for a detailed treatment, see Caro^13^). For narrow stripes, such functions would be severely limited by poor lion and hyaena acuity: only conspicuous to those predators at close range and when static. Consistent with this assumption, close observation of predator-prey interactions has not yielded clear evidence that lions or hyaenas are confused, dazzled or misled by zebras’ pelage any more than that of other ungulates^6,11,13^.

Broad rump stripes, however, are visible from greater distance: what antipredator benefits might they confer? We may confidently exclude any role in crypsis: their width and high contrast makes them highly counter-productive in that regard, being visible from distance. Similarly, any role in aposematism seems unlikely: neither lions nor leopards seem to avoid zebra, and spotted hyaena may find the size of zebra more of a deterrent than any markings. Our view, uniquely, explains why salient rump stripes might be associated with presence of hyaenas (and possibly, combinations of predators), rather than striping on the head, neck and torso.

Rump stripes are, of course, not the only salient stripes of particular salience on zebra. First, many plains zebra have large, salient stripes on their flanks, though these seem likely to perform similar functions to the rump stripes. In particular, as these are typically extensions of rump stripes, it seems likely they will act to draw attention effectively from the head and neck when they are foveated by a predator, via orientation-selective attention, then encourage attention ‘slippage’ toward the rear via object-based attention. There are also often thicker stripes at the base of the neck that are only discernible at closer range – these may draw predator attention away from the muzzle and throat. Given limited effective range and being occluded from view in fleeing zebra, their impact on pursuit per se is likely limited.

A limitation of our view, and likely a reason it has not been expounded before, is that Grevy’s zebra do not have salient rump-stripes. Their narrow rump stripes may maximize protection from tabanids, but provide no benefit of cueing predator attention to the rear. Perhaps, as for the African wild ass and extinct quagga, minimising conspicuity at a distance may confer compensatory benefits in small or heterospecific groups^24^. Alternatively, the Grevy’s markings may reflect reduced selective pressure from lion and hyaena predation relative to plains zebra^25^. Reduced adaptation might also contribute to reportedly disproportionate selection of Grevy’s lions and spotted hyaenas^26,27^.

We have assumed here that zebra stripes evolved primarily to deter ectoparasites, and that the antipredator function described here essentially piggy-backed on that mechanism. However, there remains controversy over zebra stripes’ primary function^28^. Some studies have concluded that temperature, rather than parasites, were primary drivers of stripe evolution, including rump stripes^8^. Such findings may yet implicate thermoregulation as zebra stripes’ primary function; it is certainly a key variable in predator-prey interactions, though currently evident for cooling effects of zebra stripes is lacking^15^. Moreover, that view (as with other extant accounts) does not explain for the striking differences in widths of rump stripes versus other stripes, and the correlation between temperature and stripe patterning breaks down if mountain zebra are also considered.

Our findings account for a puzzling aspect of zebra stripes by estimating their appearance through the eyes of predators. Our conclusions do not rest on the accuracy of these estimates or assumptions about salience. Under the conditions specified, any visual system with human acuity or worse, irrespective of the shape of the predator’s contrast-sensitivity function, should yield the same effects. This mechanism, we can be confident, should exert a strong exogenous (stimulus driven) pull on predators’ attention during pursuits, conferring clear advantages to zebra.

## 3 Methods

### 3.1 Resource Availability

#### Lead Contact

Further information and requests for resources should be directed to and will be fulfilled by the Lead Contact, Greg Davis.

#### Materials Availability

Visual stimuli used in the study will be deposited on GitHub: https://github.com/alexmuhl-r/Zebra-Project.

#### Data and Code Availability

The datasets and code generated during this study will be made available on GitHub (URL TBC).

### 3.2 Experimental Model and Subject Details

#### Observers

Observers between 18 and 45 years old with normal or corrected-to-normal vision and access to a desktop or laptop computer were recruited via Prolific.co. Observers in Experiment 1 were automatically precluded from participating in Experiment 2. Twenty-four observers completed Experiment 1 (*M* _age_ = 23.75 years, *SD*_age_ = 3.78, 16 females, 8 males, 4 observers were excluded and subsequently replaced because of missing mouse tracking data). Twenty-eight observers completed Experiment 2 (*M* _age_ = 24.46 years, *SD*_age_= 6.26, 9 females, 19 males, 1 observer was excluded and subsequently replaced because of missing mouse tracking data). These sample sizes were selected to ensure that we could detect effects of size *F* = 0.25 in our planned ANOVAs and of *d* = 0.5 in our planned t-tests with 80% power. These sample sizes and power calculations were pre-registered (for further information on pre-registration see Method Details below). These experiments were approved by the Cambridge Psychology Research Ethics Committee at the University of Cambridge and informed consent was obtained from all observers.

### 3.3 Method Details

#### Stimuli

Forty-eight photographic images of single zebra against naturalistic backgrounds were used as visual stimuli in Experiments 1 and 2. These comprised of twelve images of each of the following Stripe-Patterns: Mountain Zebra, Plains Zebra (no shadow stripes), Plains Zebra (shadow stripes), Grévy’s Zebra. The images were drawn from online photograph repositories and were selected to show an unobstructed view of the zebra in profile. Using the GNU Image Manipulation Program, three images were edited to remove highly visually salient genitalia and then all images were cropped to fit closely to the zebra. A horizontal motion blur with a length of 50 px was applied to produce an additional set of 48 motion-blurred images. All 96 of these images were then scaled down to 960 px in width (original aspect ratios preserved) and then placed on a 1024 px x 1024 px square Gaussian noise background.

#### Experiment 1

For Experiment 1, these images with a Gaussian noise background were then scaled down to 512 px x 512 px. All images were adjusted to simulate the photopic visual acuity of lions using the AcuityView package in R^29^. This adjustment assumed a viewing distance of 70 cm, image width of 13 cm and a visual acuity of 13.42 cpd21. Further, all images were then colour adjusted to simulate lion/hyaena dichromatic vision using Peter Kovesi’s colour blindness simulation script in MATLAB26^30^ with peak sensitivity values of 430 nm for the short wavelength cones and 553 nm for the long wavelength cones^21^. Finally, every image was flipped horizontally to create both a left- and right-facing version. The resulted in a final set of 196 stimuli (512 px x 512 px) that were used in Experiment 1, all of which were colour- and acuity-adjusted, including simulated motion and static images and left-facing and right-facing images. Simulated motion and static stimuli were always presented in separate order-counterbalanced blocks.We used a screen calibration procedure (see Salience Task section below) to ensure that these images were always presented as 14 cm x 14 cm irrespective of differences in display resolution (a small deviation from the value used in our initial acuity adjustment, but consistent across all stimuli in Experiment 1). At a size of 14 cm x 14 cm, these stimuli had a visual angle of 2.86° at a viewing distance of 70 cm and 4.01° at a viewing distance of 50 cm. Within this plausible range of viewing distances and assuming an average real-world zebra length of 2.42 m (averaged values from Caro^13^), these stimuli produce a retinal image the same size as viewing a zebra at between 8.64 m and 12.10 m.

#### Experiment 2

For Experiment 2, the 48 non-filtered 1024 px x 1024 px Gaussian noise background images were scaled down to produce a set of 256 px x 256 px images and a set of 165 px x 165 px images (images in this latter set were all placed on an additional Gaussian noise background to increase the overall image sizes to 256 px x 256 px so that they were compatible with the AcuityView package). All images were adjusted to simulate three different visual acuity levels to simulate the photopic acuity of lions (13.42 cpd), an average of the mesopic acuity of lions and photopic acuity of hyaenas (7.84 cpd), and the mesopic acuity of hyaenas (4.60 cpd) using the AcuityView package in R. These adjustments assumed a viewing distance of 70 cm and an image width of 7 cm (the image width parameter for the 165 px x 165 px images was not smaller than the 256 px x 256 px images because of the additional Gaussian noise border added). All images were then colour adjusted to simulate lion/hyaena dichromatic vision as for Experiment 1. For each Striping Pattern, at each acuity level and image size, half of the images were randomly selected to be flipped horizontally for one group of observers and the other half for a second group of observers, resulting in a counterbalanced and equal number of left- and right-facing zebra for each group of observers. This resulted in a final set of 288 stimuli that were used in Experiment 2, at two sizes (165 px x 165 px and 256 px x 256 px) and three acuity levels (lions photopic, lions mesopic/hyaena photopic, and hyaena mesopic). As in Experiment 1, we used a screen calibration procedure to ensure that he 165 px x 165 px images were always presented as 3.5 cm x 3.5 cm and the 256 px x 256 px images as 7 cm x 7 cm irrespective of differences in display resolution. At a size of 7 cm x 7 cm, the 256 px x 256 px stimuli had a visual angle of 5.72° at a viewing distance of 70 cm and 8.01° at a viewing distance of 50 cm. Within range of viewing distances and assuming the same average real-world zebra length of 2.42 m, these stimuli produce a retinal image the same size as viewing a zebra at between 17.28 m and 24.22 m. At a size of 3.5 cm x 3.5 cm, the 165 px x 165 px stimuli had a visual angle of 11.42° at a viewing distance of 70 cm and 15.94° at a viewing distance of 50 cm. With the same assumptions as above, these stimuli produce a retinal image the same size as viewing a zebra at between 34.56 m and 48.47 m.

#### Salience Task

The salience task was an online, browser-based, task developed and hosted using the Gorilla Experiment Builder^31^. The purpose of this task was to obtain estimates of the most visually salient locations on our zebra stimuli. We used mouse click location to gather these estimates by instructing observers to click on the part of the image that caught their attention. This approach provided a significantly higher level of precision that presently available webcam eye tracking technology. Observers completed the task on their own computer in a location of their choosing and responded with mouse clicks. In addition to only recruiting observers with access to a desktop or laptop computer, observers were prevented from beginning the task if they were not using a web browser on a desktop or laptop computer. On each trial of this task, observers were shown a central fixation cross which they were instructed to click when they were ready to start the trial. Three hundred milliseconds after this click an image of a single zebra appeared in the centre of the screen (see Stimuli section) and observers were instructed to look at the zebra and then to click on the area of the zebra that caught their attention. Observers were told not to reflect on this but rather to just click wherever ‘caught their eye’. Mouse movements were recorded during each trial and, following a mouse click on the zebra, a blank screen was shown for one second before the next trial began. Observers received breaks of at least ten seconds every 48 trials. The task would be automatically terminated and observers excluded if not completed within two hours of starting to protect against drop out. Additional Javascript code was used to check that the task was always run in fullscreen mode and a screen calibration procedure, using a credit or debit card as a reference object, was used at the beginning of the task to ensure that images were presented at a specific size regardless of observers’ specific computer hardware. While this allowed the dimensions in pixels of each image to vary, it ensured that images were a consistent physical size on all screens irrespective of small variations in display resolution. Observers were instructed to sit approximately 50 cm from their computer screen, while this figure did match the viewing distance used when acuity-adjusting our stimuli, we expected that most observers would not be able to sit 70 cm from their computer screens and this was also consistent across all stimuli. As an additional confirmation of image size, at the beginning and end of the task, observers were asked to click on the corners of a reference image that was the same size as the zebra stimuli and these coordinates were recorded.

#### Experiments 1 and 2 Procedure

Experiment 1 had 196 trials with motion-blurred and non-blurred stimuli presented in two order-counterbalanced blocks to which observers were randomly assigned (trial order within blocks was randomised). Experiment 2 had 288 trials in randomised order with two counterbalanced sets of stimuli (different halves of the stimuli facing different directions) to which observers were randomly assigned. Beyond these differences, both experiments followed the procedure outlined below. After signing up on Prolific.co^32^ observers were redirected to our task hosted via the Gorilla Experiment Builder where they initially viewed an information sheet and, if they wished to take part, completed a consent form. If they consented, observers were directed to the salience task, which initially checked observers were viewing the task in fullscreen mode, observers could only continue by clicking a button which activated fullscreen mode if it was not already active. Following this check, observers completed a screen calibration procedure, using a credit or debit card as a reference object, to ensure that images were presented at a specific size regardless of observers’ specific computer hardware. At this point, observers were instructed to ensure they had a stable internet connection before continuing further. Observers were then told that the task would involve viewing images of zebra and that they would need to click on these images while their mouse movements were recorded. Before more specific instructions, observers completed an additional confirmation of image size, by clicking on each of the four corners of a reference image that was the same size as the zebra stimuli (this procedure was also repeated at the end of the task). Observers were then given details instructions for the main part of the task. For each trial, observers were instructed to take a moment to look at the zebra and then click on the area of the zebra that caught their attention. Breaks of at least ten seconds were included every 48 trials. After the task observers were presented with a debriefing which explained the aims of the experiments and highlighted some relevant literature, they were also encouraged to report, by typing into a text box, any particular parts of zebra to which their attention was drawn.

### 3.4 Computation Modelling

#### Bottom-up salience maps

To estimate bottom-up visual salience, salience maps were generated using the SMILER software implementation^33^ in MATLAB R2020a. This package allows the generation of 14 different computational instantiations of bottom-up salience maps. While feature map parameters can be individually specified, for simplicity, maps were generated using default parameters. Salience maps were generated for all right-facing stimuli and were identical to those seen by observers. No additional transformations were performed. For the model comparisons with human responses, we used the LDS model (Learning Discriminative Subspaces)^34^, as this model resulted in tightly distributed peaks in salience similar to those made by human observers when clicking on the stimuli. These maps allowed analysis of the association between human and model salience estimates, using Pearson correlations of the two-dimensional salience distributions. We report human-model correlations for two additional salience models, the Graph-based Visual Salience (GBVS)^35^ and Fast and Efficient Saliency (FES)^36^ models. These correlations are reported in Supplementary Tables 1 and 2.

#### Predator-prey pursuit simulations

This simulation estimated the time to intercept for a predator (e.g., lion or hyaena) pursuing a zebra when the predator tracking the front versus the rear of the zebra, at several starting distances, and predator top speeds. The simulation was programmed in MATLAB R2020a. The simulation initialised with the zebra and predator taking positions (*x,y*) in a featureless landscape. The positions (*x,y*) of the predator and the zebra were advanced iteratively, such that the zebra moved in a straight line in *x*, while the predator heading altered based on the angle between itself and the front or rear of the zebra. Both predator and prey were initially static and accelerated at a constant rate until reaching their respective top speeds. The simulation ended when any one of the following criteria was met: 1) if the predator *x* coordinate exceeded the rearmost point of the zebra, indicating successful interception; 2) if the rate of change of the difference between the zebra and predator positions in *x* increased, indicating that the zebra has outrun the predator; 3) if the chase time exceeded 20 s, indicating that the predator might have ceased pursuit. When the simulation ended, the number of iterations was recorded and converted to a ‘time-to-capture’, expressed in seconds. The simulation made several simplifying assumptions. Firstly, the model was updated in one millisecond iterations. While it is not feasible that a predator could update its heading on the basis of visual feedback in such short intervals, it enabled finer-grained estimation of the difference in chase time than using a larger iterative interval and is unlikely to have affected the outcome of the simulation. Secondly, there was no delay between the initiation of movement for the predator and zebra. In the real-world, both predator and prey will demonstrate short delays in responding to visual information. It is also likely that one animal will move before the other, with either the predator initiating a chase and prey then fleeing or the prey fleeing at the sight of the predator. Thirdly, the terrain is featureless, with no obstacles, cover or elevations. Finally, both animals accelerate at a constant rate and the zebra moves in a straight line, without responding to the heading of the predator to maximise distance or engage in evasive, protean, behaviours. Complexities introduced by cooperative hunting are also ignored, though note that benefits in the modelled scenario effectively augment the range of angles at which zebra can flee, disrupting predator strategies of corralling prey toward cooperating predators. The simulation has several fixed parameters and several experimentally manipulated parameters. Zebra body length was fixed at 2.42 m with an initial position centred on the origin and the zebra was represented by a line between points (−1.22, 0) and (1.22, 0). The zebra had a fixed top speed of 17 m/s and a constant acceleration of 5 m/s2. The predator had an initial position perpendicular to the centre of the zebra (*x* = 0) and a *y* coordinate that was manipulated to determine predator-zebra distance at the beginning of the chase (minimum starting distance 10 m, increased in 5 m increments to a maximum of 55 m). The predator had a constant acceleration of 9.5 m/s^2^ and its top speed was manipulated between the values of 14 m/s and 20 m/s in increments of 1 m/s. Both predator and prey began to move on the first iteration of the simulation. The critical manipulation was whether the predator aimed for the front or rear of the zebra. The front was operationalised as forward-most point of the zebra the rear was operationalised as its rearmost point.

#### Pre-registration

Experiments 1 and 2 were pre-registered at AsPredicted.org (Experiment 1 registered 25/11/2020, AsPredicted.Org #52944; Experiment 2 registered 16/12/2020, AsPredicted.Org #54384).

#### Quantification and Statistical Analysis

Statistical analysis was carried out in R and MATLAB. The results and statistical details of all analyses can be found in the Results section. A standard alpha level of *–* = 0.05 was used. Sample sizes were selected to ensure that we could detect effects of size *F* = 0.25 in our planned ANOVAs and of *d* = 0.5 in our planned t-tests with 80% power (these sample sizes and power calculations were included in our pre-registration). Trial order randomisation was achieved using built-in functions in the Gorilla Experiment Builder.

#### Experiments 1 and 2 Analysis

The data from Experiments 1 and 2 were analysed in R and these analyses were pre-registered (see above). No data were excluded on the basis of our pre-registered criteria for excluding the data of any observer who clicked on a single side of the stimuli for >74% of trials or who demonstrated a bias for a single side of the stimuli at least 2.5 standard deviations from the sample mean for any zebra species. The dependent variable of interest for Experiments 1 and 2 was the proportion of clicks on the front vs the rear of the zebra. We calculated proportion scores for each observer and stripe-pattern, subtracting 0.5 from each proportion to centre the range at zero. For Experiment 1, we then conducted four separate 2 x 2 repeated measures ANOVAs (motion blur: (Static, Simulated-Motion) x facing: left, right), one for each species. We conducted two follow-up 2 x 2 x 2 ANOVAs which included species as two additional factors ([Grevy’s, Plains without shadow stripes], [Grevy’s, Mountain]). For Experiment 2, to simplify our analysis, the direction stimuli were facing was not included as a factor in our analysis and we conducted four separate 3 x 2 repeated measures ANOVAs (acuity: lion photopic, hybrid lion/hyaena, hyaena mesopic x size: middle-distance, far), one for each species. We expected to see, in intercept term, evidence for bias > 0 (towards rear) for plains zebras without shadow stripes and for mountain zebras. We expect the effect to be weaker/absent for the other two types of zebra. We also conducted pairwise t-tests to follow up the ANOVA results for Experiment 2.

#### Computational Modelling Analysis

As expected, the model indicated that irrespective of predator speed and the starting distance between the predator and prey, aiming at the rear of the zebra rather than the front increases the time-to-capture, and therefore the likelihood that the predator will catch the prey. In fact, in the case that a predator has a lower top speed than their prey (but a greater acceleration), there will be starting distances at which aiming at the rear rather than the front will be decisive in whether the predator catches the prey or does not, with rear-aiming predators failing to catch their prey, even in cases where the entirety of the chase is over within 5 s for the front-aiming predator (hatched areas; Figure 4, right panel). It should be noted that as the simulation makes several necessary simplifying assumptions regarding the evolution of the predator-prey pursuit, it is unlikely that the projected times-to-capture would be correct for real pursuits. However, withstanding idiosyncratic pursuit characteristics, the pattern of results might be expected to generalise across predator-prey pursuits for which there are similar speed and acceleration differences.

## 4 Acknowledgements

We thank Alexandra Woolgar, Rebecca Smith, Kayleigh Paske, Daniele Campello, of University of Cambridge, for assistance sourcing images. A second, parallel project on human targeting of striped images involved an initial experiment with Jennifer Daffron and subsequent work with Rory Durham and Josh Lloyd. Completion of that work has been delayed by the COVID-19 pandemic.

## 5 Author Contributions

AM and MGP contributed equally to experimental design, stimulus preparation, data collection, computational modelling, pursuit modelling, statistical analysis, and manuscript preparation. GD conceived of the project and contributed to experimental design and manuscript preparation.

---

**Supplementary Table 1.** Mean Pearson correlation coefficients between computational model salience and observers’ salience estimates (Experiment 1)

|  | LDS |  | FES |  | GBVS |  |
| --- | --- | --- | --- | --- | --- | --- |
|  | SIM | Static | SIM | Static | SIM | Static |
| Grevy's | .378 | .183 | .503 | .350 | .291 | .298 |
| **Mountain** | **.419** | **.212** | **.479** | **.271** | **.318** | **.262** |
| **Plains-** | **.457** | **.363** | **.543** | **.552** | **.311** | **.411** |
| Plains+ | .477 | .280 | .510 | .505 | .295 | .332 |

**Supplementary Table 2.**
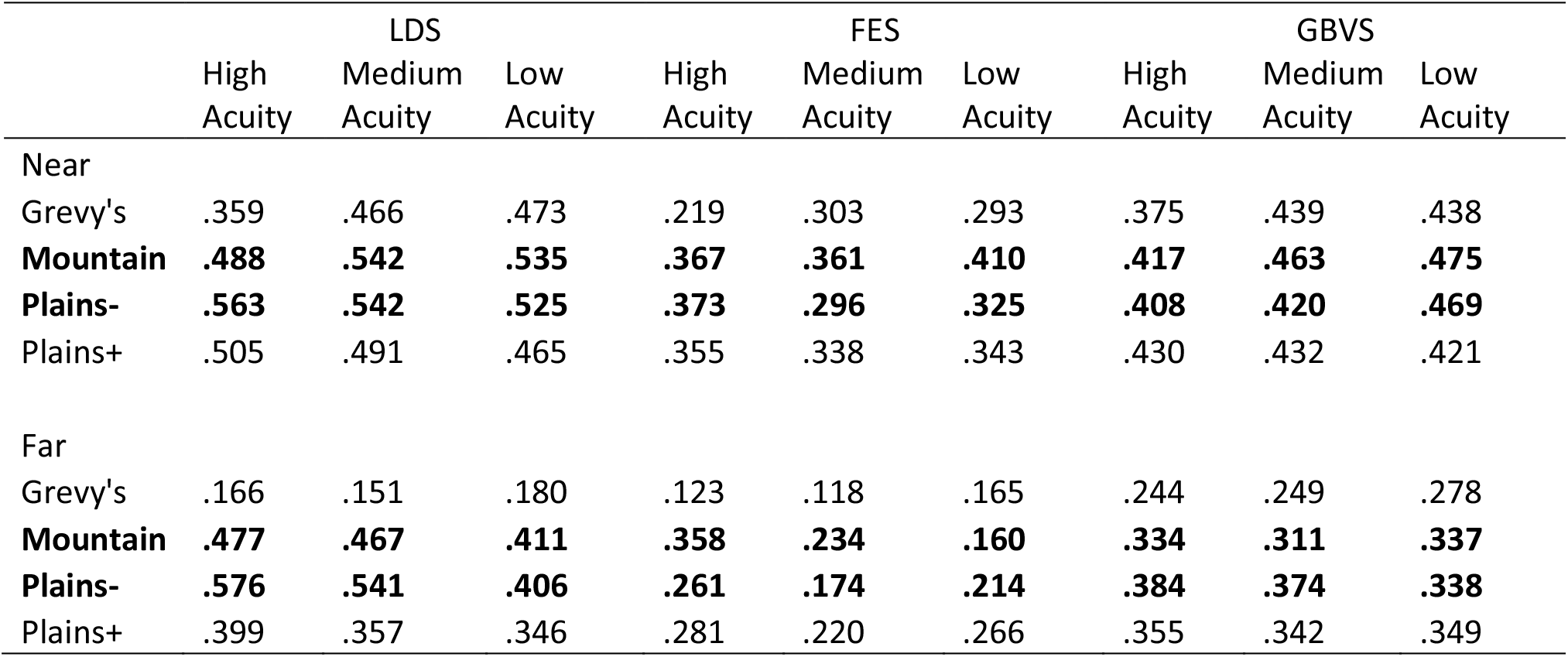
Mean Pearson correlation coefficients between computational model salience and observers’ salience estimates (Experiment 2)

